# Climate Warming Patterns Predict Potential Residency Shifts

**DOI:** 10.64898/2026.09.11.750976

**Authors:** Gabriel Hessler, Shilin Ma, Neil Gilbert, Donald DeAngelis, Lu Zhai, Bo Zhang

**Author notes:** Corresponding author: Bo Zhang, ^1^Department of Biology, Oklahoma State University, Stillwater, OK, USA, 74074.

## Abstract

**Introduction and Aim:** Future climate instability creates obstacles for organisms across taxa, especially in areas that experience higher stochasticity. Wetland wading bird populations are a prime example of this scenario, as the hydrologic cycles that drive population success in these habitats depend heavily on the predictability of wet and dry periods. When fluctuations in wet and dry periods shift excessively, populations may be forced to relocate or shift migration routes. Our goal is to predict if worsening environmental conditions could drive increases in facultative migration and subsequent residency shifts

**Location:** State of Florida, USA.

**Methods:** We developed an individual-based movement model motivated by wading birds in southern Florida to determine if this concept of climate-driven range or residency shift is likely to occur in the future. To further interpret the outputs of this individual-based model, we compared habitat suitability for the American wood stork across Southeast of the United States between 2010 and 2025, 2055 and 2085

**Results:** Our IBM model predicted that a northward residency shift is predicted to occur given the environmental scenario of harsher fluctuation via climate instability in the Everglades. Moreover, our findings suggest that wood storks with partial and/or facultative migration may be more likely to adapt to these fluctuations. Our suitability maps showed spatially heterogeneous responses, with notable increases in northern Georgia, and the Carolinas, and consistent declines in northern Florida, indicating a gradual northward shift in suitable breeding habitat.

**Main Conclusions:** Conserving networks of suitable wetlands across broader spatial scales, may be critical for facilitating movement and persistence under hydrologic instability. Mechanistic models such as the one presented in this study can help anticipate these shifts and inform proactive conservation planning. This model can also be adapted to fit similar wetland systems, or other landscapes with various population and movement phenotypes present.

## Introduction

Climate warming and concomitant increases in extreme weather events threaten ecosystem stability and biodiversity (Larsen et al. 2011). Other forms of human disturbance can amplify these effects of climate change. (Bouton et al. 2005; Butler and Vennesland 2000). Wetland ecosystems are at particular risk because they are highly sensitive to the wet–dry cycles imposed by local climate conditions (Xiong et al. 2023; Canning and Waltham 2021). Small wetlands are often heterogeneous in their capacity to support foraging across seasonal cycles. Nevertheless, many migratory species depend on these patches as stepping-stones to larger, more stable habitats (Szangolies et al. 2022; Saura et al. 2014; Rocha et al. 2021; Ramos et al. 2020) and some species even use them year-round to exploit high prey densities during drying periods (Lee et al. 2025; Yurek et al. 2024). Under climate change, many wetlands are expected to experience alterations in hydrologic regimes, leading to population declines, local extinctions, or range shifts among wetland-dependent species (Sandhu et al. 2016; Xu et al. 2024; Frederick et al. 2009).

A critical question for conserving biodiversity in light of global change is whether migrating animals can adjust their migration routes and schedules to track key resources as resource phenology shifts with climate change (Teitelbaum et al. 2016; Thorup et al. 2007). Movement is an energy investment that can be optimized for the specific environment the organism inhabits. For instance, migratory birds track food resources that occur along their routes, a process that influences their body condition and survival (Thorup et al. 2007). These food resources, which are needed for refueling and completion of the migratory journey (Linscott and Senner 2021), can undergo dramatic seasonal and inter-annual fluctuations (Tomotani et al. 2018). If stochastic variation increases in these environments, some organisms may exhibit adaptive movement strategies to increase fitness (Winkler et al. 2014; Martin et al. 2015; Ruokolainen et al. 2011). Facultative migration exemplifies these adaptive movement strategies. Facultative migration occurs when a species migrates only under extenuating circumstances such as failure of key food resources (Widick et al. 2025). With increasing environmental stochasticity, facultative migration may also increase and drive range shifts (Picardi et al. 2020; 2022; Holt et al. 2022; Butler and Vennesland 2000; Larsen TH et al. 2011) which can create further ramifications, including increased invasion risk (Wallingford et al. 2020; Brodie et al. 2025; Hodgson et al. 2022). However, it remains largely unknown when, where, and under what conditions facultative migration will emerge in response to increasing environmental stochasticity, and whether individual-level movement strategies can be used to predict which populations are most likely to undergo such shifts.

The American wood stork (*Mycteria americana*) offers an ideal system to study the impact of facultative migration on range shifts because they are a facultative migrant that uses dynamic wetland habitats. In the United States, the wood stork occupies the Everglades of South Florida for at least part of its life cycle, but ranges as far north as South Carolina, and as far west as Louisiana (see Supplementary Fig. 1). Understanding their migration decisions and assessing whether facultative migration benefits survival have important conservation implications. Southern Florida’s hydrologic cycle varies from year to year, and fluctuations of the drying down of water levels in the breeding season, when prey must become concentrated for reproductive success, can be especially critical (Stys et al. 2017). Changes in weather in the future could lead to more hydrologic variation, as the frequencies of droughts and heavy rainfall events are expected to increase (Wang et al. 2014; Beasley 2023; Abiy et al. 2019). Generally, wood storks will linger in their permanent residency year-round if they can, unless prey density begins to decline or competition between other wading birds becomes too intense (Picardi et al. 2020; Yurek et al. 2024; Gawlik 2002). In this scenario, they may take the option of long-distance migration to the north (Picardi et al. 2020). Alternatively, some storks may find secure feeding areas or adequate habitats along their migration routes and remember these locations for future use (Lee et al. 2022; Bell 1990). Using knowledge of the distribution of wetland habitats, we can provide context when observing behavioral changes across this population for current environmental fluctuations. However, it remains unknown how the distribution and reliability of wetland foraging sites for wood storks will change under climate warming and increasing hydrologic instability, and how these changes will influence individual migration decisions and range shifts, but plausible scenarios can be simulated.

Our goal is to predict if worsening environmental conditions could drive increases in facultative migration and subsequent residency shifts. We first developed an individual-based modeling (IBM) approach that tracks animal movement and environmental conditions over long timescales and incorporates inter-individual variation in movement behavior (Grimm and Railsback 2005). In contrast to existing models that primarily focus on local-scale movement and foraging patterns (Lee et al. 2022; DeAngelis et al. 2021), competitive behaviors (Yurek et al. 2024), and other environmental cues such as predation (Bracis and Wirsing 2021; Carter and Finn 1999), our framework introduces a novel individual-based model that integrates multiple movement modes of the American wood stork. Within this individual-based model we also tested how the selectiveness of individuals affects survival in this stochastic framework. To further validate the outputs of this individual-based model, we obtained raster datasets representing habitat suitability for the American wood stork across Florida, Mississippi, Alabama, Georgia, and South Carolina. We aim to answer two central questions: (1) Will birds shift from permanent southern residence to more northern areas under environments with more hydrologic fluctuation? (2) Does selectiveness have an impact on survival and/or spatial distribution? (3) Does facultative migration provide resistance to habitat instability?

## Methods

Methods based on the individual-based model used here are organized using the Overview, Design Concepts, Details (ODD) protocol (Grimm et al. 2020) created to efficiently delineate complex theoretical models and all the parts that make these models. Simulations were executed in MATLAB version 2021b and data analyses were executed in RStudio (version 4.2.2) and Excel. Additional models such as the habitat suitability rasters were executed in RStudio (version 4.5.1) and ArcGIS.

### Overview

#### Purpose

Wading bird population dynamics, movement, behavior, and other life-history facets have been studied by both empirical field studies and theoretical models (Picardi et al. 2020; 2022; DeAngelis et al. 2021; Yurek et al. 2024; Lee et al. 2025; Basille et al. 2020), using a similar IBM or agent-based modeling approach as used in this study. Our model is intended to elucidate how sequences of movement decisions over broader timescales, especially under changing climate conditions, scale up to influence species distributions.

#### Entities, state variables, and scales

There are two entities in this model, individual adult birds and patches of potentially usable habitat. The quantity of individuals can be changed; however, 200 individuals are used in a 100 × 1000 km landscape containing 360 spatially explicit patches (Fig. 1). Individuals are independent entities and, although they may cross paths throughout the simulation and even share patches, co-occurrence does not influence individual behavior, but competition can occur when more than one wading bird occupies a site. 360 patches are randomly distributed along the x-axis and are separated into three geographic regions along the y-axis: south (0-200 km), central (201-799 km), and north (800-1000 km), roughly reaching a region north from southern Florida. The southern and northern regions contain 140 patches each, and the central region contains 80. This configuration was chosen for the present study, but the framework is readily extensible and can be modified to accommodate other systems, spatial arrangements, or research objectives.

**Figure 1:**
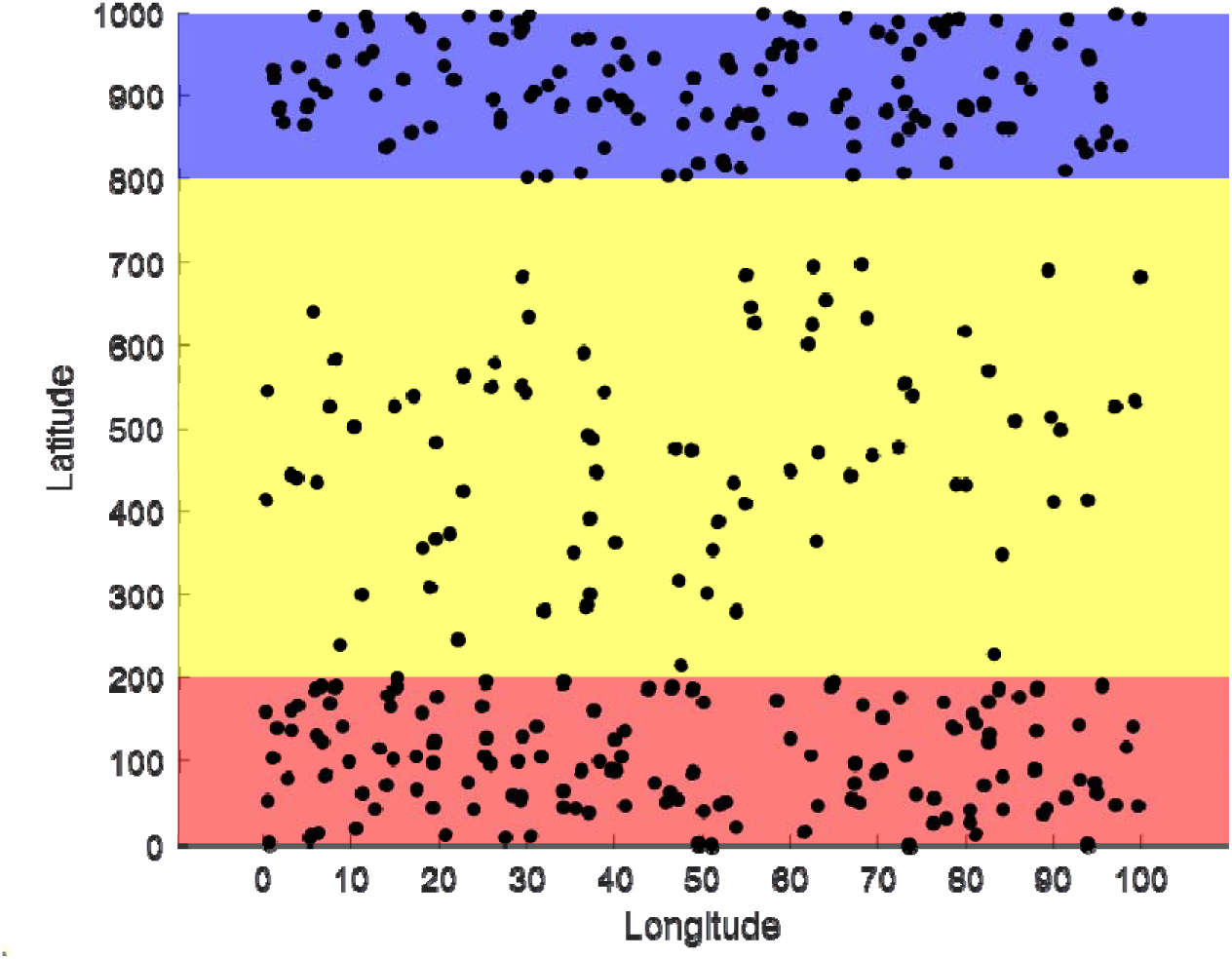
Spatial distribution of 360 patches across the 100 × 1000 km simulated landscape. Patches are split into three regions:140 in the south (red region), 80 in the central yellow region, and 140 in the north (blue region). These divisions are abstract representations of sites in southern, central, and northern Florida.

#### Process overview and scheduling

All simulations take place over a 20-year period in daily timesteps (7305 days) for each prey density threshold (10 total). Each patch can change sizes (effecting foraging favorability) each timestep based upon environmentally relevant scenarios, with larger patches containing more food. Patches cannot go extinct and instead can reach a minimum size that offers exceedingly low food even for a single individual. Within each prey density threshold, the model follows wading birds for 1 day where the birds survey the landscape for one suitable patch for that timestep. This is repeated for all 200 wading birds and energy intake is calculated (as described in *Energy Parameters* below). Birds with no energy die, and then the simulation moves to the next day of movement. This process is repeated for 7305 daily timesteps within each of the 10 prey density thresholds. This simulation is repeated for 10 replicates for each environmental scenario. The order of the simulation is as follows: prey density threshold begins (20-40) Timestep begins (1-7305) Bird movement (1-200 birds) Energy calculations Timestep ends prey density threshold ends. See figure 2 for a schematic of the bird movement loop and pathing.

**Figure 2:**
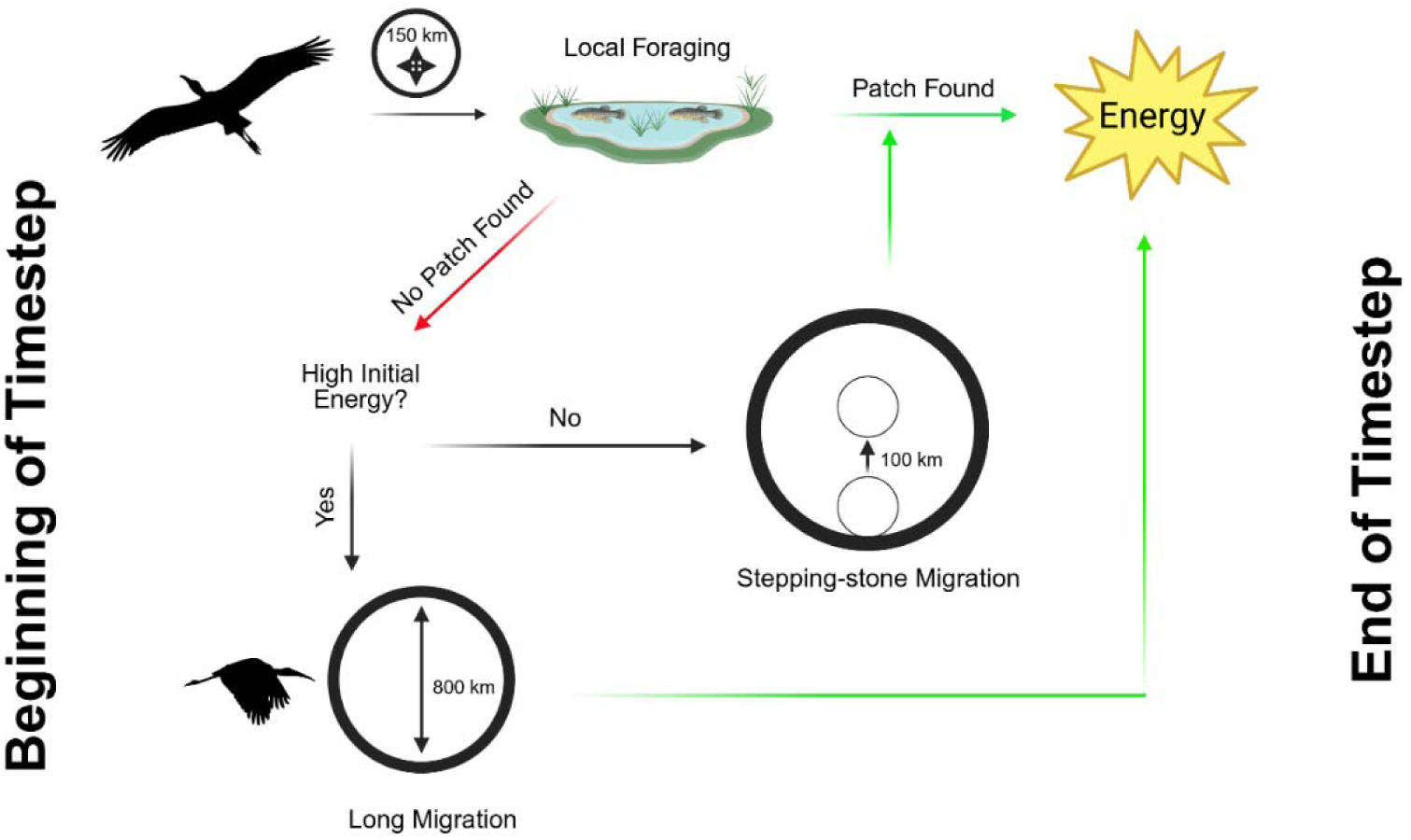
Bird movement path that occurs during each timestep for each bird. Once the energy icon is reached, the loop begins again with a new bird.

#### Prey Density Thresholds

Multiple prey density thresholds mimic selectiveness in foraging behavior, indicating if higher or lower preference for prey density, alters survival or final location. There are 10 prey density thresholds that dictate decision making of an individual. These prey density thresholds are values that correlate to the minimum prey density that a bird will tolerate within a patch, mimicking selectiveness, and are directly influenced by patch size. For example, at prey density threshold 1, birds must land on a patch size greater than or equal to 20. As prey density threshold increases up to level 10, the minimum patch size that birds land on increases up to a size equal or greater than 40, which is exceedingly rare during given environmental scenarios. These thresholds are equal for all birds during simulations.

### Design Concepts

#### Emergence

There are 3 emergent properties in the model. The first is patch selection by local foraging and stepping-stone foraging within a single day. The second is movement strategy if local foraging attempts fail which provides the choice between stepping-stone and long-migration across the landscape. The third is spatial distribution of individuals across the landscape at the beginning and end of any period of time.

#### Adaptation

There is only one adaptation available to individuals. This adaptation is movement strategy, where two additional movement options (long-migration and stepping-stone migration) are available if a bird fails in its local patch foraging method.

#### Objectives

The objective of an individual is to survive daily metabolic burn by foraging within a suitable patch and retrieving energy.

#### Learning

There is no direct or indirect learning within the simulation.

#### Aggregation

Although wood storks are social and often migrate in flocks, we chose not to explicitly model aggregation behavior to preserve parsimony.

#### Sensing

Birds can sense/survey up to 30 patches within a set radius of 150 km that can be slightly altered by whatever movement strategy they utilize during a timestep. The presence of other birds is not assumed to be sensed; that is, wading birds are not using others as cues to either visit or avoid a patch.

#### Stochasticity

Individual patch sizes fluctuate along a pre-set range where size can fall anywhere within that range during any given timestep. For the southern patches in the experimental environment, this fluctuation is biased with time, where the maximum and minimum size of a patch decreases as time increases (Fig. 3A). For the control environment, patch sizes fluctuate equally regardless of time or region (Fig. 3B).

**Figure 3:**
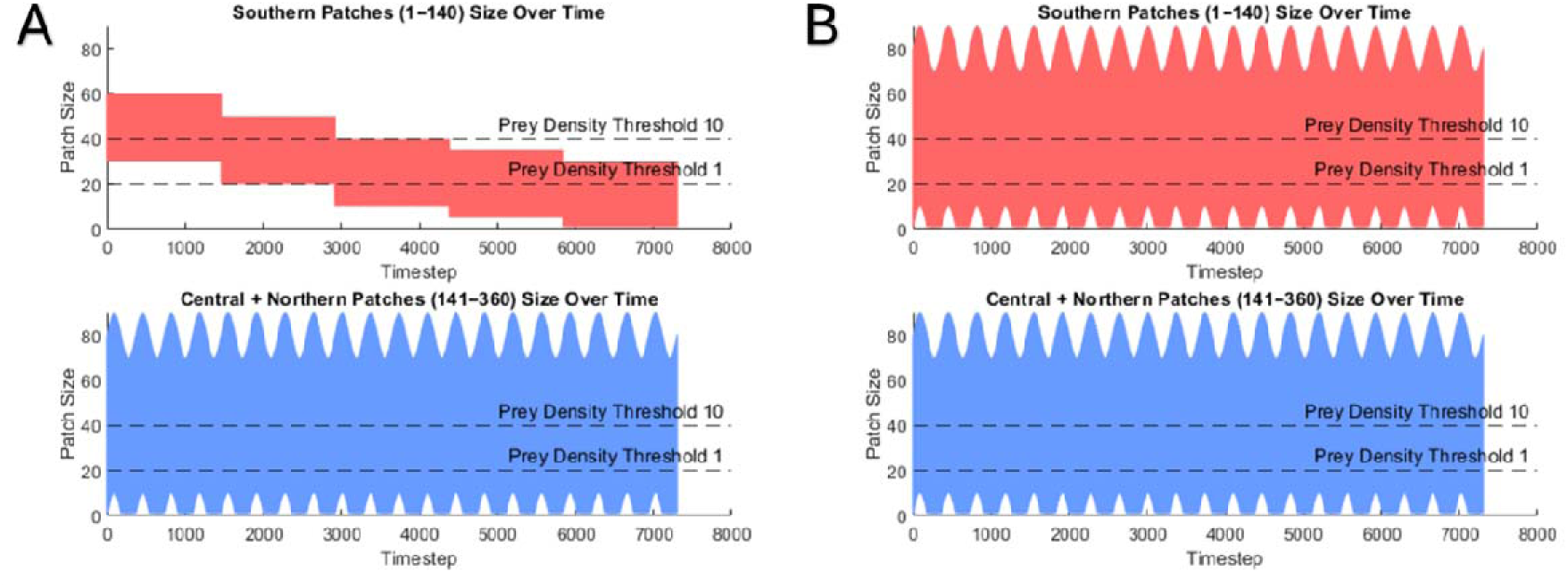
Simulated environment patch size ranges. (A) Patch size fluctuations in the experimental environment with degrading conditions in the southern region only. (B) Patch size fluctuations in the control environment with equal fluctuations across all regions. Gridlines at 20 and 40 are included to show the maximum and minimum prey density thresholds (1 and 10).

#### Parameterization

Parameters for both the individual and environment were based on wood stork information pulled from observational field studies and other models (Wolff 1994; Blake 1982; Kushlan 1979; J. M. Hefner, “Wetlands of Florida, 1950s to 1970s,” 1986) Environmental parameters have been scaled from the shape and structure of the state of Florida (approximate distance from the Everglades to northern border). Patch size fluctuations were created using expert opinion from members of the United States Geological Survey (USGS) in Everglades National Park.

#### Observations

The one major point of data collection within the model simulation for this project is the location of individuals throughout time, since the overarching goal of this project is to identify factors that can lead to a shift in permanent residence. Apart from changing location (longitude + latitude), we also collected several other points of data throughout the simulation, some of which are mainly collected to run certain functions in the model itself but can also be indicators for foraging success. For example, energy via prey consumption was collected throughout the entirety of the simulation, and average prey consumption in certain parts of the environment can indicate suitability of that environment. Mortality events were also collected and identify in what region and conditions led to the death of a bird.

### Details-Initialization

#### Starting locations and conditions for individuals

At time T_1_, each bird is assigned to a random patch within the southern region or northern region of the landscape depending on the present scenario (see *Stochasticity* above). Each bird starts with 900 calories of energy, which is double the daily metabolic rate of wild adult wood storks (Wolff 1994; Kahl 1964) Birds do not move during T_1_, instead they collect the energy at their current patch and end the timestep. Competition is not considered for T_1_.

### Details-Submodels

#### Energy Parameters

Initial energy at T_1_ for all birds is 900, which is double the daily metabolic burn of the American wood stork. After T_1_, the initial energy of a bird is the energy gain calculated at the end of the previous timestep. Energy calculations vary based on the movement option used during the timestep. For the default method (30 local patches) and stepping-stone method, energy at the end of the timestep is calculated as (Energy_Gain_ = Energy_Initial_ + Food_Patch_ – Metabolic_Burn_). Birds do not feed during long-distance migration, so energy at the end of the timestep is (Energy_Gain_ = Energy_Initial_ – Metabolic_Burn_). For the fallback option where no movement options worked, energy is calculated as (Energy_Gain_ = Energy_Initial_ – Food_Patch_ (Distance_Penalty_) – Metabolic_Burn_). If the distance traveled is greater than 400 km but less than 800 km, Distance_Penalty_ = 0.5. This cuts the energy gained from the patch by half. If distance traveled is greater than or equal to 800 km, Distance_Penalty_ = 0 to completely negate Food_Patch_ in the fallback equation. If distance traveled is less than 400 km, Distance_Penalty_ = 1 to add no consequence to food gained. Whenever a bird’s energy declines to 0, the bird dies and is removed from that prey density threshold iteration after its location and movement strategy is saved. Initial energy or energy gained cannot exceed 1800 (4 times the metabolic burn).

#### Competition

To properly model resources shared if a patch is occupied by several birds, we use a simple penalty added to “Food_Patch_” where we divide the food at a patch equally among the number of individuals at that patch. This competition penalty is not applied to T_1_ to avoid immediate death.

#### Scenarios

Four scenarios are described in this model across two environments. Our experimental environment models a wetlands landscape with climate change altering hydrologic cycle in the south, while the central and northern regions fluctuate along the same range independent of time (see *Stochasticity* above for further details). Our control environment models a wetlands landscape with equal patch fluctuation throughout all regions. Using these environments, the 4 scenarios used in this study are as follows: (1) Experimental environment with all 200 birds starting each prey density threshold in the southern region. (2) Experimental environment with 100 birds starting in the south, and 100 birds starting in the north. (3) Control environment with 200 birds starting in the south. (4) Control environment with 100 birds starting in the south, 100 birds starting in the north.

#### Data analysis

A total of ten replicates were completed for each of the 4 scenarios which includes the 10 prey density thresholds per replicate. Mean-latitude of survivors and mortality rate within each prey density threshold for each replicate was collected and exported to Excel. We used a one-way ANOVA to test the significance of prey density threshold on mortality and mean-latitude within each scenario (*R Core Team 2021*). To test the significance of each scenario, and to compare the results of scenarios against one another, we used a two-way ANOVA with Tukey post-hoc pairwise comparisons. Conclusions from these analyses are dispersed throughout the results section when applicable.

### Habitat Suitability Tests

To further interpret the outputs of this individual-based model, we obtained raster datasets representing habitat suitability for the American wood stork across Florida, Mississippi, Alabama, Georgia, and South Carolina. These datasets were developed by the National Audubon Society and include 1-km resolution suitability surfaces and associated derivative layers produced from species distribution models for 604 North American bird species. Projections are available for a recent distribution (2010) and for two future climate change scenarios (RCP 4.5 and RCP 8.5) for the 2025s, 2055s, and 2085s. Suitability values ranging from 0-1 were multiplied by 10,000 for better representation of differences. To estimate temporal changes in wood stork suitability, we computed difference rasters by subtracting the baseline (2010) suitability surface from each projected layer. This calculation was performed separately for breeding-season and non-breeding-season suitability, and for both climate scenarios (RCP 4.5 and RCP 8.5) across all three future time horizons. The resulting rasters represent the magnitude and direction of suitability change relative to the 2010 baseline. All raster processing and visualization were conducted in R 4.5.1 using the *terra* (Hijmans 2025) and *ggplot2* packages (Wickham 2016). For each of the 12 suitability-change rasters, we extracted minimum and maximum cell values to characterize the full range of projected change. We then identified the global minimum and maximum across all rasters to standardize the color scale used for mapping. To enable direct comparison among scenarios and time periods, we applied a unified diverging color scheme in which positive values (increased suitability) were displayed in red, negative values (decreased suitability) in blue, and zero-change values in neutral gray. Cells with missing data (NA) were rendered in white to clearly distinguish areas without suitability information.

## Results

The IBM simulations examined how prey density thresholds—representing levels of foraging selectivity—influence wading birds’ selection of foraging sites, alongside the use of different movement and foraging strategies. There were four scenarios, with and without climate change in each of the two simulated landscapes. Overall, our simulation results showed that higher prey density thresholds were associated with higher mortality in every scenario (p < 0.001). Indeed, high prey density threshold was also associated with higher mean-latitude of survivors in the experimental landscape (scenarios 1 and 2, p < 0.001) but not in the control landscape (scenarios 3 and 4). The initial spatial distribution of birds did not have a significant effect. This is especially noticeable in visual comparisons between scenarios, as the results from scenario 1 are extremely similar in most replicates to the results from scenario 2. The same trend is carried over to scenarios 3 and 4. To simplify visual aids because of that trend, scenarios 2 and 4 have been moved to the supplementary material.

### Southern degradation over time increases mortality

When the southern region of the landscape degraded over the 20-year timespan, we observed high mortality rates and northward residency shifts. In both scenarios 1 and 2 with different initial spatial distribution of birds, mortality exceeded 15% at low prey density thresholds and increased further, peaking at over 50% as prey density thresholds increased to 10 (Fig. 4A). This was a linear curve with higher prey density thresholds equating to higher mortality (Fig. 4A), but there was no significant difference in mortality rates between scenarios 1 and 2. Consequently, more northward permanent residencies of survivors (green dots in middle panel of Fig. 1a) were observed for scenarios 1 and 2, regardless of prey density threshold. More importantly, most survivors occurred in the northern region at a prey density threshold of 1, while at a threshold of 10 the southern region had no survivors and the highest mortality. The mean-latitude of surviving birds was significantly higher in scenario 2 compared to scenario 1 (p = 0.04), indicating a benefit to birds who start the simulation in the more stable northern region.

**Figure 4:**
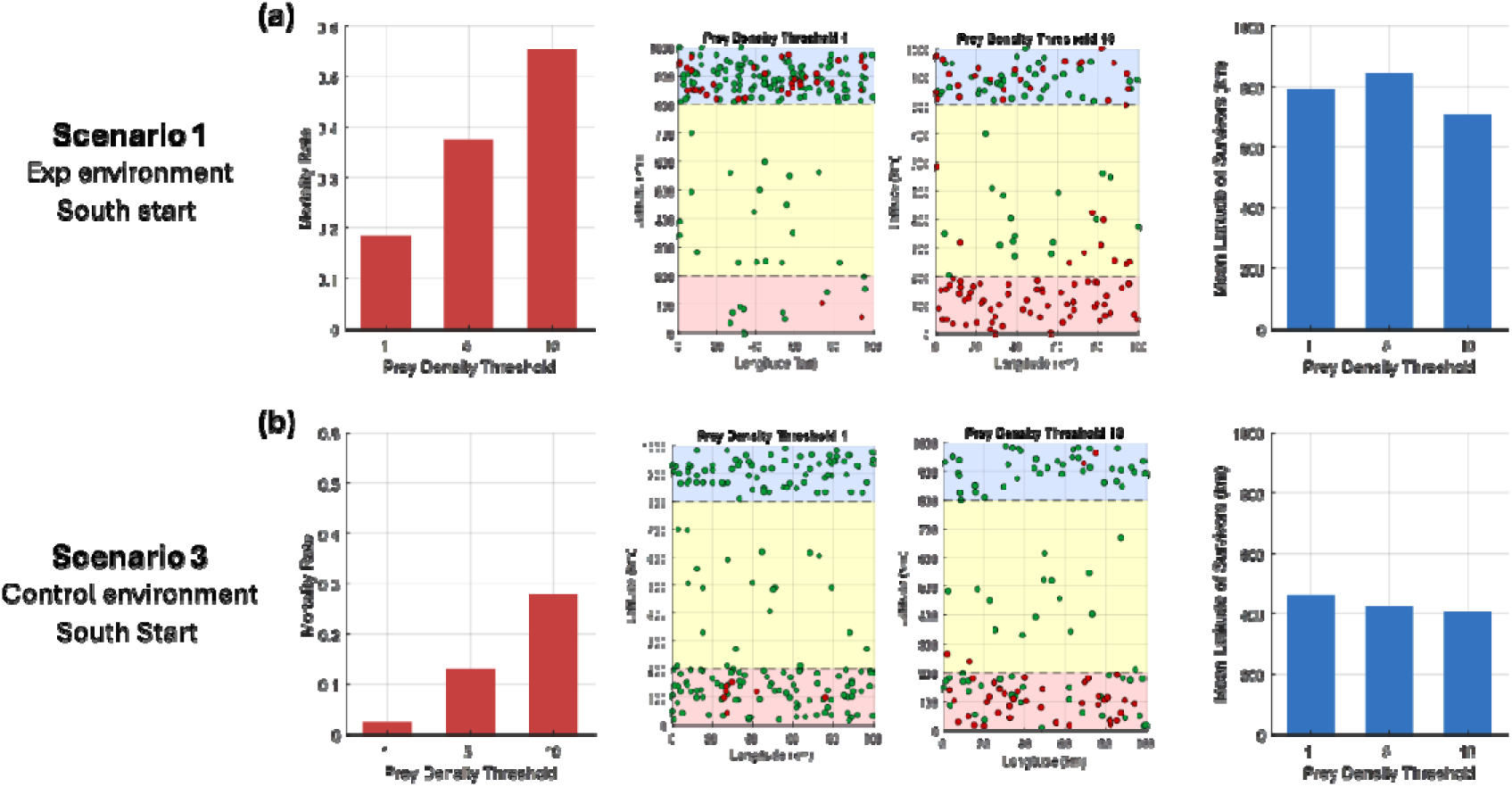
An example of simulation outputs for scenarios 1 (A) and 3 (B). The left panel shows mortality rates across prey density thresholds 1, 5, and 10. The middle panels show spatial distribution of surviving birds (green dots) and dead birds (red dots) across the landscape in prey density thresholds 1 and 10. The right panel shows the average latitude of surviving birds in prey density thresholds 1, 5, and 10. All panels show data after the full 20 year timespan. See S2 for visualizations of scenarios 2 and 4.

### Equal landscape fluctuation provides stability (scenarios 3 and 4)

In scenarios 3 and 4, which have equal fluctuation throughout all regions, mortality rates were generally lower, and the latitude of survivors was more varied. Mortality rates were rarely above 0-3% for the first few thresholds across 20 years, especially in scenario 4, which had significantly lower mortality rates overall, regardless of threshold, compared to scenario 3 (p < 0.001). However, both scenarios 3 and 4 had significantly lower mortality compared to scenarios 1 and 2 (p < 0.001 for all, Fig. 4B), indicating that equal landscape fluctuation across all regions provides more stability. Spatial distribution of surviving birds is more uniformly distributed across the entire landscape, rather than being solely located on one region, as in scenarios 1 and 2 (Fig. 4B). This trend is also transferred to the distribution of dead birds, even in higher prey density thresholds. The mean-latitude of survivors was generally located between 300-600 for every replicate regardless of prey density threshold, with no significant difference between scenarios 3 and 4. Much like mortality however, mean-latitude was significantly lower in scenarios 3 and 4 compared to scenarios 1 and 2 respectively (p < 0.001 for all).

### Facultative migration provides adaptation to stochasticity

When the southern region of the landscape degraded over the 20-year timespan, northward residency shifts were facilitated. Under these conditions, both long-distance and stepping-stone migration strategies were expressed. Migration frequency for both strategies exhibited a positive linear increase through time (Fig. 5A–B), with stronger increasing trends observed at higher prey density thresholds. Alternatively, long-distance and stepping-stone migration was less common in the control environment and did not follow any linear trend as seen in the experimental environments (Fig. 5C-D), suggesting that stable landscape conditions limit the expression of migratory strategies.

**Figure 5:**
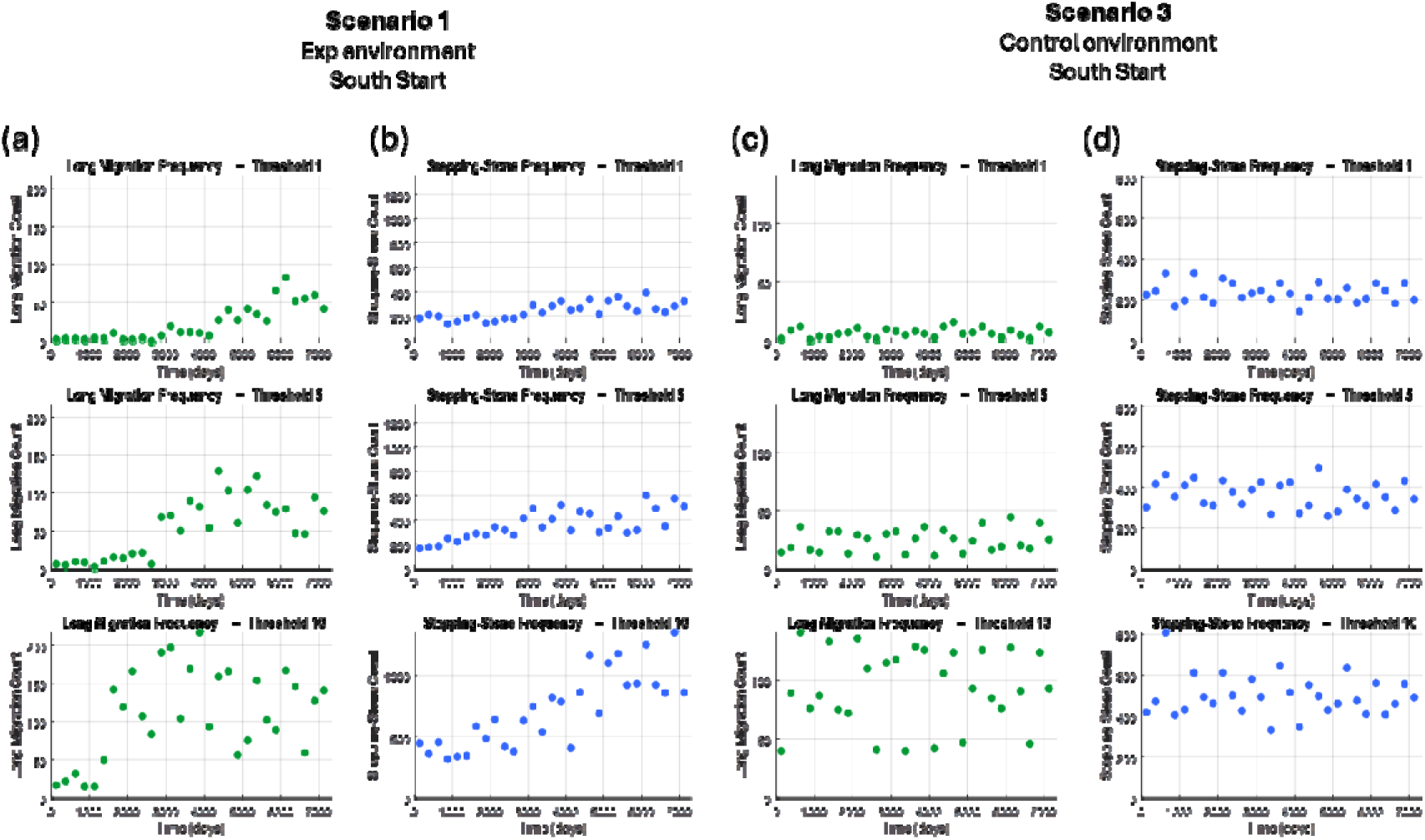
Frequency of long-distance and stepping-stone migration in scenarios 1 and 3. Each dot notes the total events of that migration type every 250 timesteps across the entire 7305 timestep simulation. (A) Long-distance migration frequency in scenario 1. (B) Stepping-stone migration frequency in scenario 1. (C) Long-distance migration frequency in scenario 3. (D) Stepping-stone migration frequency in scenario 3.

### Habitat suitability maps suggest stability in the north

As obtained from the habitat suitability tests, across both seasons, suitability generally increased over large portions of the region, with changes becoming more pronounced over time and under the higher-emissions scenario. Projections under breeding season showed spatially heterogeneous responses, with notable increases in northern Georgia, and the Carolinas, and consistent declines in northern Florida and parts of the Gulf Coast, indicating a gradual northward shift in suitable breeding habitat (Fig. 6). In contrast, projections under nonbreeding season exhibited more widespread positive changes, with most of the Southeast showing increasing suitability and only limited declines in southern Florida. Overall, the maps suggest an expansion of climatically suitable conditions in northern areas and a contraction at the southernmost range edge, with RCP 8.5 producing the largest magnitude of change.

**Figure 6:**
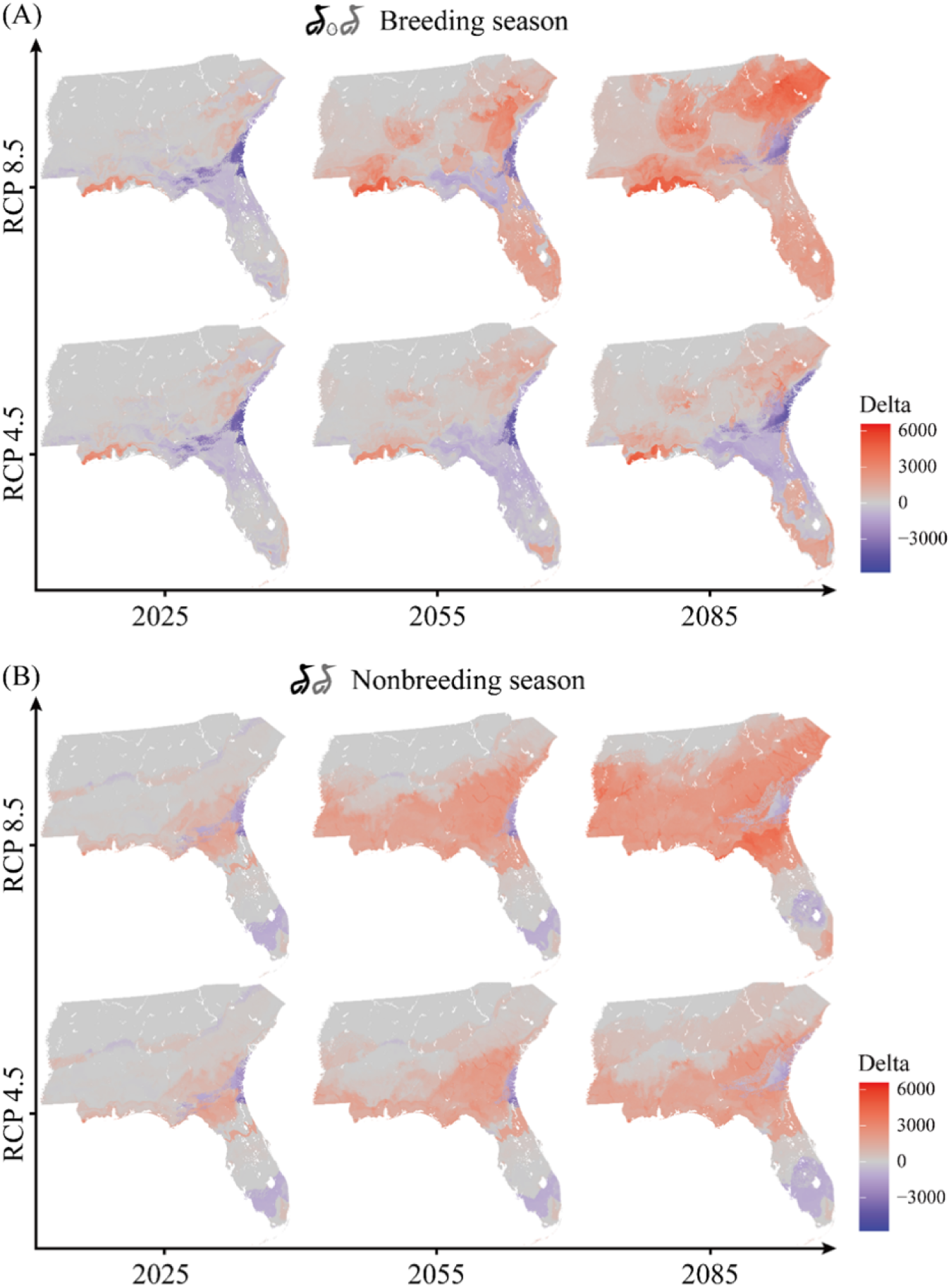
Projected changes in wood stork suitability across the southeastern United States for both the breeding season (A) and nonbreeding season (B), under two climate change scenarios (RCP 4.5 and RCP 8.5) and three future time periods (2025s, 2055s, and 2085s). All values represent deviations from the 2010 baseline, with positive changes shown in red (increased suitability), negative changes in blue (decreased suitability), and near-zero changes in gray. Suitability values ranging from 0-1 were multiplied by 10,000 for better representation of differences.

## Discussion

Our individual-based model indicated that increasing hydrologic instability in south Florida can drive emergent range shifts. Across simulations, degradation of southern patches led to increased mortality, greater reliance on facultative migration, and a redistribution of survivors towards northern regions of the landscape. The interpretation(s) of the IBM results are further strengthened given the context of the habitat suitability maps and potential instability of south Florida in the future. If the northern regions become significantly more suitable across breeding and non-breeding seasons, it is likely that a residency shift could occur for this population.

### Facultative migration as a response to hydrologic instability

Under experimental scenarios where southern patch quality declined over time, facultative migration became increasingly common, especially when prey density thresholds increased (Fig. 5A-B). Both long-distance and stepping-stone migration strategies were utilized when local foraging failed and often led to higher survival if it meant an individual escaped the deteriorating south. In fact, increased use of these migration strategies coincided with higher survival in northern regions, suggesting that facultative migration can function as an adaptive mechanism against local habitat degradation, supporting theories from similar studies (Gilroy et al. 2016; McCrary et al. 2019). In contrast, when patch quality fluctuated evenly across all regions in the control environment, migratory strategies were used less frequently and showed no consistent temporal trend (Fig. 5C–D). Mortality remained low across prey density thresholds, and survivors were more evenly distributed across the landscape. This contrast highlights that migration in the model is not intrinsically advantageous but becomes adaptive only under sustained spatial asymmetry in habitat quality. The context that this trend provides can further improve our understanding of migration evolution (Salewski and Bruderer 2007; Bruderer and Salewski 2008; Zink 2011). The patterns in this model suggest that increasing climatic variability, rather than average conditions alone, may play a critical role in shaping movement strategies and residency outcomes.

### Selectiveness influences outcomes

Variation in prey density thresholds strongly influenced both mortality and spatial distribution across scenarios, highlighting the effect of selectiveness in dynamic environments such as wetlands. Higher selectiveness equated to higher mortality as suitable patches became exceedingly rarer under southern degradation and general fluctuation in the control environment (Fig. 3). Selectiveness had a linear trend with spatial distribution in the experimental environment only, where higher selectiveness pushed birds north. It is important to re-iterate that birds are all given the same level of selectiveness depending on the whatever level of prey density threshold the simulation is running. It may be unlikely in nature to see this occur. What may be more realistic is a gradient of selectiveness across a population with individuals being able to actively make selective decisions based upon social (Dickie and Serrouya 2022) and environmental cues (Lyu et al. 2021). Based upon the results of the individual based model, we would expect similar trends, however, with more selective individuals finding more success in stable northern regions as the south possibly degrades over the next century.

### Independent habitat suitability projections support directional consistency

While the individual-based model does not explicitly incorporate climate variables, the habitat suitability projections provide independent spatial context that is consistent with the emergent patterns observed in the simulations. Across climate scenarios and time horizons, suitability generally declined in southern Florida while increasing at higher latitudes, particularly in the southeastern United States (Fig. 6). These projected shifts mirror the northward redistribution of survivors produced by the model, suggesting that climate-driven changes in habitat quality may reinforce the behavioral mechanisms identified here. Importantly, the habitat suitability analysis is not intended as a validation of the individual-based model, nor does it capture the mechanistic processes underlying movement decisions. Instead, it provides a complementary, pattern-based perspective that converges on the same directional outcome. The agreement between mechanistic simulations and correlative projections strengthens confidence in the broader inference that climate instability may promote northward residency shifts in facultatively migratory wetland species.

### Limitations & Future Directions

As with any modeling study, the results from this model are subject to several limitations. First, seasonal differences, that is breeding vs. non-breeding, are not explicitly considered to reduce complexity over the 20-year timespan. Second, learning and evolution are not involved in simulations, once again to reduce complexity. Third, while social dynamics such as competition and aggregation are present in the model, they are heavily simplified. Lastly, patch and environmental dynamics in the individual based model are influenced mostly by expert opinion rather than actual climate data. We would expect that the addition of these factors would show similar outcomes in the end, perhaps with even greater selection for northward shift. The difficulty comes in accurately incorporating these factors into the model to improve realism rather than introduce confounds.

### Conservation Implications

From a conservation perspective, our findings suggest that species exhibiting facultative or partial migration strategies may possess greater defense against increasing environmental variability and/or instability. This defense, however, may come at the cost of increased mortality, especially depending on the selectiveness of individuals. Additionally, if populations migrate away from historical residencies, then the new residences the population will then occupy may have positive or negative effects on the existing ecosystem of those new residences. Conserving networks of suitable wetlands across broader spatial scales, including potential stepping-stones, may be critical for facilitating movement and persistence under hydrologic instability. Mechanistic models such as the one presented in this study can help anticipate these shifts and inform proactive conservation planning. This model can also be adapted to fit similar wetland systems, or other landscapes with various population and movement phenotypes present.

## Acknowledgements

1. G. H., S.M., L.Z, and B.Z. acknowledge financial support from the NSF Mathematical Biology program (2325196). We’d like to thank Simeon Yurek and Simona Picardi for their assistance in the early design phase of this study.

## Biosketch

Gabriel Hessler is a researcher studying movement ecology, with a focus on how animals navigate spatially complex landscapes. His work combines theoretical and experimental modeling in the laboratory to investigate how environmental conditions, resource distribution, and movement strategies shape population abundance and persistence.

## Funding Statement

G. H., S.M., L.Z, and B.Z. acknowledge financial support from the NSF Mathematical Biology program (2325196).

## Conflicts of Interest

The authors declare no conflicts of interest.

## Authorship Contributions

G.H., N.G., B.Z., and D.A. gathered background information and designed this study. G.H., D.A., and B.Z. created the individual-based model. S.M. and L.Z. created the habitat suitability maps. G.H., N.G., D.A., S.M., and B.Z., provided constructive feedback and contributed to the final manuscript.

## Data availability

Information regarding the historical range and spread of *Mycteria americana* were derived from cited sources, with range map permissions from South Dakota Birds and Birding (https://www.sdakotabirds.com/index.html). Data and code involved with the IBM results are publicly available online (https://doi.org/10.5281/zenodo.4026064). Data and layers used for the habitat suitability modeling are also publicly available through Audobon (<u>Audubon</u> <u>Climate-Based Bird Distribution Models for North America | AdaptWest</u>).

## Ethics Statement

The authors have nothing to report.

## Notes

https://doi.org/10.5281/zenodo.4026064

### Competing Interest Statement

The authors have declared no competing interest.

