## Supplementary Figures for "Climate Warming Patterns Predict Potential Residency Shifts"

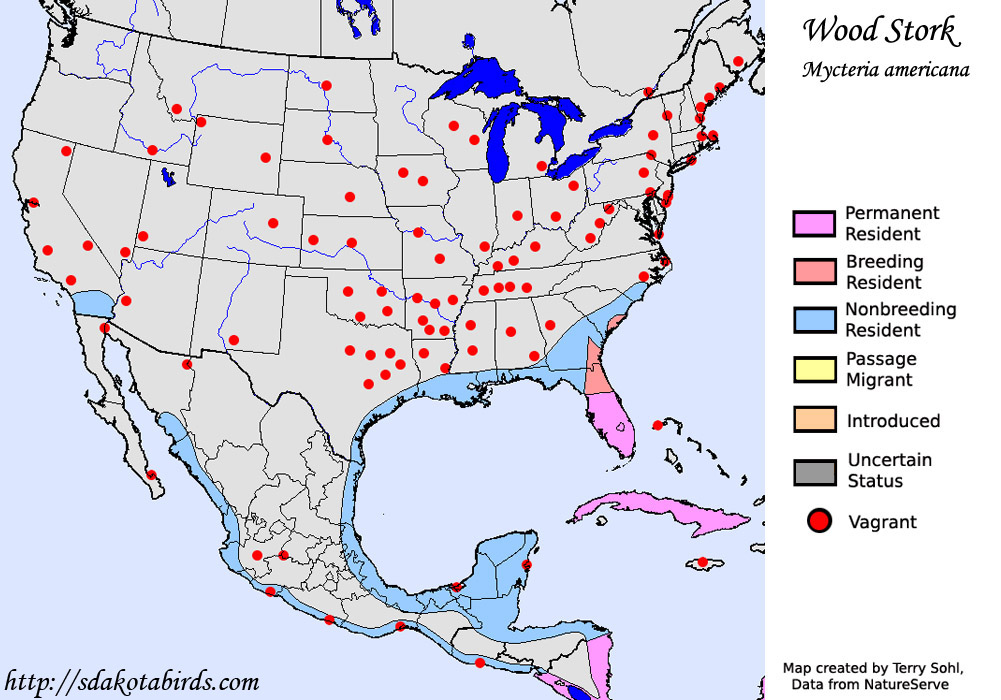

Supplementary Figure 1: Historical range map of Mycteria americana (American wood stork) in North America.
